# Mapping synaptic ensembles through *in vitro* functional cell assemblies

**DOI:** 10.1101/2024.10.10.617167

**Authors:** Clara Zaccaria, Asiye Malkoç, Ilya Auslender, Yasaman Heydari, Marco Canossa, Beatrice Vignoli, Lorenzo Pavesi

## Abstract

In memory circuits, synaptic engrams consist of ensembles of strengthened synapses that form between engram neurons encoding distinct memory traces. As proposed in Hebb’s seminal postulate, these potentiated synapses enable the functional cell assembly essential for neuronal wiring. While existing technologies can track potentiated synapses in behaving animals, understanding the *in vivo* spatial-temporal organization of synaptic engrams remains challenging. We developed an *in vitro* hybrid system that integrates digital light processing with optogenetics and used optical stimulation to synchronize the firing patterns of two-neuron modules within a network, supporting Hebb’s postulate experimentally. After illumination, we observed that synapses predominantly strengthen on dendrites between the two illuminated neurons, reflecting neuron responses to the wiring activity. This observation provides a strong conceptual foundation for our methodology. Our setup also allows for direct illumination and strengthening of synaptic ensembles, granting spatial-temporal control over the synaptic networks. Building on Hebb’s theoretical framework, our system offers a solid experimental approach to test the fundamental principles underlying synaptic engrams.

## Introduction

The term “engram” refers to the physical changes in the brain that are linked to memory processing, representing a network of neurons that form a memory trace^1–5^. The prevailing view is that the formation of an engram involves the strengthening of synaptic connections among selected neurons during memory encoding, resulting in ensembles of potentiated synapses termed “synaptic engram”^6–8^. This increased synaptic strength is thought to enhance the probability of recreating the same neural activity pattern during memory retrieval. Recent technological advancements have enabled the tracking of synapses associated with engram circuits throughout the brain during memory processes in behaving animals^3,8–11^. However, a major challenge lies in accurately mapping the spatial and temporal distribution of functional synaptic connections. This makes it difficult to fully understand how synaptic engrams encode and transmit information across the brain. As a result, establishing a direct causal link between synaptic engrams and the neural wiring that underpins memory circuits remains a complex and unresolved issue.

Engrams can extend across multiple brain regions and involve functional connectivity among distributed neurons^5,12^; however, the foundational elements of an engram can be studied on a smaller scale. Donald Hebb theoretically formulated the principle of an engram, proposing that synaptic reinforcement occurs when two neurons, A and B, are simultaneously and intensively stimulated^13^; in essence, a functional cell assembly creates the engram^14^. Building on this theoretical framework, we combined digital light processing (DLP) and optogenetics to optically stimulate the prototypical Hebbian two-neuron modules within an *in vitro* network. Our approach incorporates SynActive (SA) technology^15^, which, using both ‘live imaging’ and ‘post hoc analysis,’ enables us to detect variations in the number of potentiated spines and their distribution within the illuminated neurons. In addition, our system allows targeted illumination of selected spines to directly strengthen specific synaptic ensembles, offering precise control over the spatial-temporal organization of the synaptic network. This enables the on-demand generation of functional connectivity between neurons.

Our integrated methodology offers the opportunity to explore foundational principles of cell assemblies within a controlled *in vitro* system, aligning with Hebb’s postulate on synaptic strengthening. This provides a robust conceptual foundation for our methodological framework.

## Results

### The DLP-based system

We developed a DLP system (Figure 1A) combined with optogenetics to simultaneously stimulate individual neurons within a network. The detailed operational principle of the DLP-based system is illustrated in Figure S1 and Document S1. To assess the reliability of our system, we tested the optical resolution and optogenetic excitation on cortical embryonic cultures transduced with the adeno-associated virus AAV9-hSyn-hChR2(H134R)-EYFP (AAV-ChR2-YFP). This viral vector enabled the selective expression of Channelrhodopsin-2 (ChR2) in most neurons within our cultures (Figure S2). The cell density was calibrated to ensure sufficient network connectivity within the field of view (FOV), as demonstrated by immunocytochemical analysis using anti-SMI312 and anti-MAP2 antibodies, which target axons and dendrites, respectively (Figure S2). Light stimulation achieves varying scales of resolution, allowing for precise illumination of individual neuronal cell bodies or specific subregions of a neuron, down to single dendritic spines (Figure S1C). Neurons were optically stimulated using a standard patterned illumination protocol (Figure 1B) known to induce long-term potentiation (LTP)^15^, a paradigm associated with synaptic engram encoding^16^. Confirmation comes from experiments that combined DLP illumination of a selected group of neurons within a network and LTP recording using a Multi-Electrode Array (MEA). We observed that patterned light stimulation successfully induced LTP in certain areas of connectivity within the neuronal network (Figures 1C and 1D).

**Figure 1.**
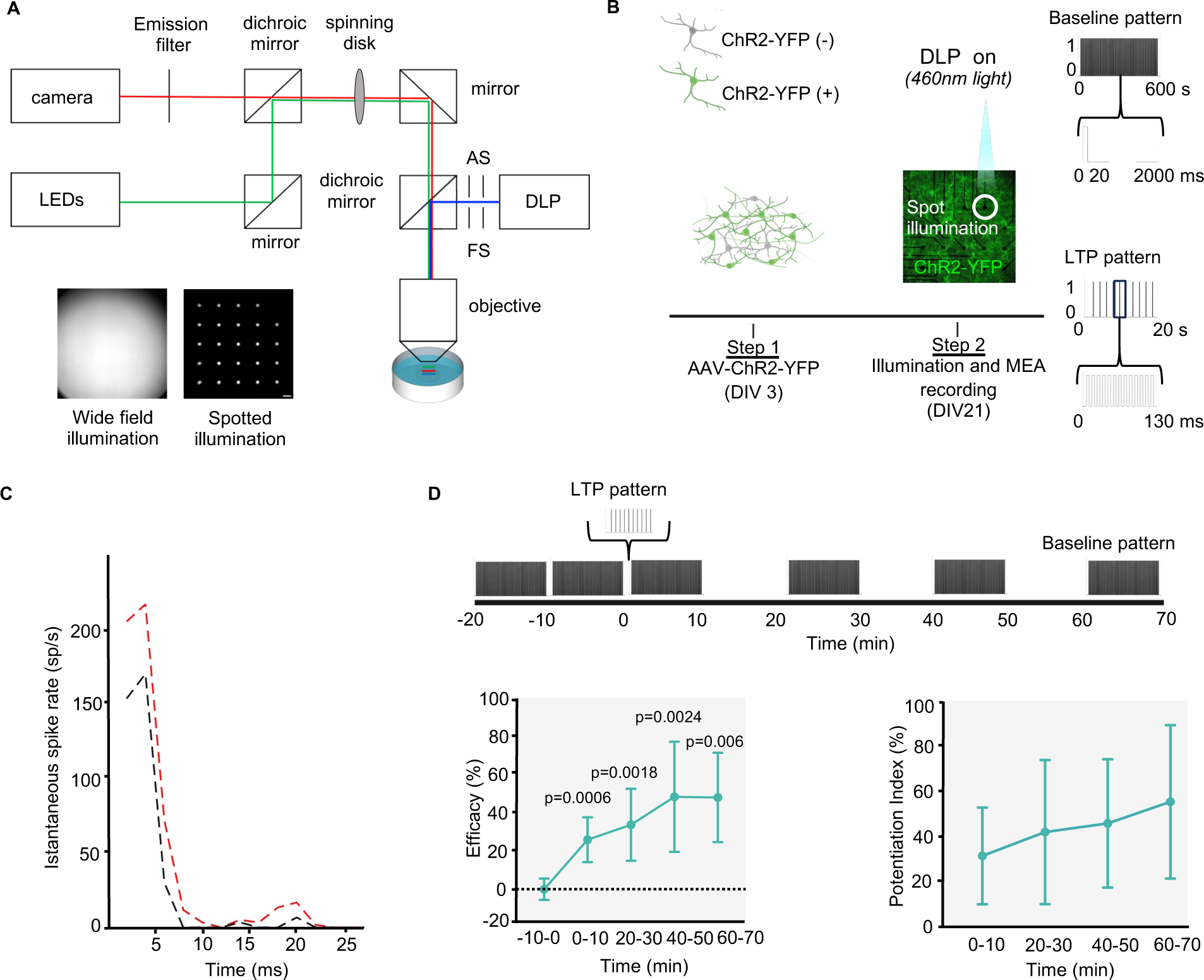
SD setup, implemented with DLP and MEA recording. (A) Scheme of the SD setup implemented with DLP. AS and FS refer to the aperture stop, and the field stop diaphragms. Representative light patterns for wide field and multiple spot illumination are shown. (B) Schematic diagram of the experimental procedures and of the temporal patterns used for baseline (20 ms light pulses at a frequency of 0.5 Hz for a duration of 10 min) and LTP light stimulation (10 sets of 13 light pulses delivered at a frequency of 100 Hz with a repetition frequency of 0.5 Hz). (C) PSTH profiles representing the responses of the network before (black) and 20 min after (red) the LTP stimulation. Bin size = 2 ms. (D) Upper, timeline representation of the MEA experimental sequence. Lower, efficacy and potentiation index measured at different time points after the LTP light stimulation (n=6 experiments; Unpaired t-test).

### Generation of prototypical cell-assembly modules

Our primary goal was to generate discrete modules of light-activated cells, serving as prototypical examples of Hebb’s “cell assemblies” for engram circuits^14^. Using patterned light on neurons A and B, simultaneously, we generate modules of two-activated neurons (Figure 2A). Initially, we assess neuron responsiveness by imaging Ca^2+^ fluctuation before, during, and after the light stimulation using the Ca^2+^ indicator X-Rhod-1 (Figure 2B). We show that most neurons of the two-neuron modules responded to optical stimulation with a significant increase in X-Rhod-1 fluorescence intensity (ΔF/F) compared to nearby non-illuminated neurons within the FOV (Figure 2C). When light stimulation was applied to individual neurons, we observed a similar increase in ΔF/F levels (Figure S3), suggesting that potential synergistic activity between the two illuminated neurons does not significantly affect the Ca^2+^ response. Moreover, variations in ChR2-YFP expression levels did not affect Ca^2+^ responsiveness (Figures 2C and S3C), indicating that variations in ChR2-YFP did not compromise the accuracy of our Ca^2+^ measurements.

**Figure 2.**
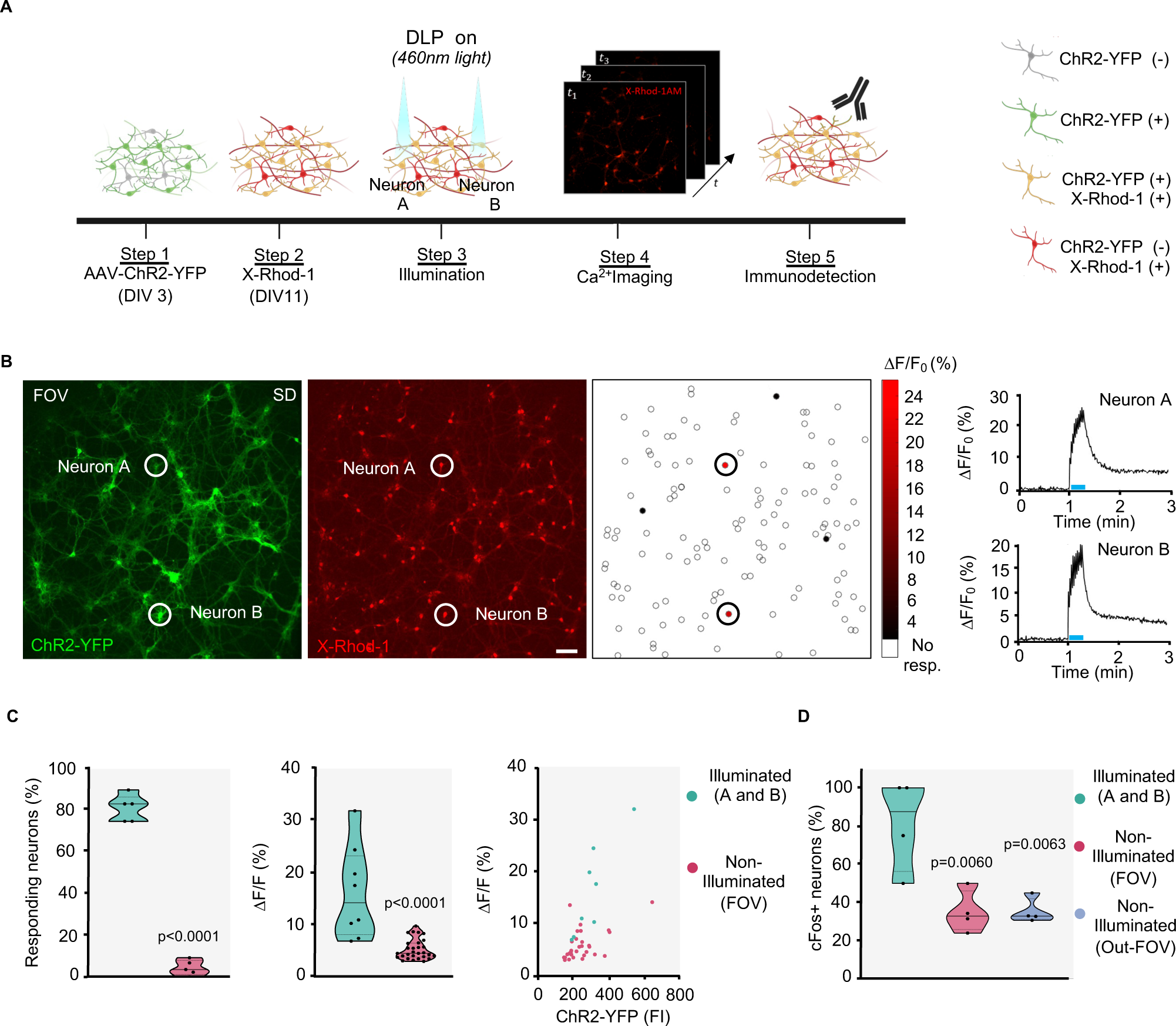
The prototypical cell-assembly module. (A) Schematic diagram of the experimental procedures. (B) SD images depicting ChR2-YFP and X-Rhod-1 fluorescence signals in a representative neuronal network. Scale bar: 50 μm. Illuminated neurons (A and B) are highlighted (circles). A heat map illustrates the percentage change in fluorescence (ΔF/F) calculated in automatically detected ROIs. The diagrams display fluorescence changes (ΔF/F) recorded from the soma of neurons A and B, before, during, and after light stimulation (blue line). (C) Left, quantification of number of responding neurons A and B and non-illuminated neurons in the FOV (n=5 experiments; Unpaired t-test). Middle, quantification of ΔF/F (n=8 cells for illuminated (A and B); n=29 for non-illuminated (FOV); Mann-Whitney test). Right, ΔF/F plotted against the ChR2-YFP fluorescence intensity (FI) (n=8 cells for illuminated (A and B); n=31 for non-illuminated (FOV). (D) Percentage of A and B neurons, non-illuminated neurons within the FOV and Out-FOV expressing cFos (n=4 experiments; One-way ANOVA Test).

Next, we examined whether light stimulation induces changes in the expression of cFos, an immediate early gene (IEG). This gene is commonly utilized as a marker to identify active engram neurons, and IEG promoters are used to recruit engram neurons in behavioral experiments^12,17^. Ninety minutes after stimulation, cFos expression was elevated in neurons exposed to the light stimulus compared to non-illuminated neurons within the FOV or outside (Figure 2D).

In summary, we integrated optical stimulation, optogenetics, electrophysiology, Ca^2+^ imaging, and marker-based activity assessments to activate prototypical two-neuron modules within a neural network.

### Spatial-temporal analysis of synaptic strengthening

To validate the functional assembly of neurons A and B in our experimental setup and to empirically support Hebb’s postulate, we investigated whether patterned light stimulation could induce synaptic strengthening between the illuminated neurons. We integrated our system with SynActive (SA) technology, a genetically engineered platform that utilizes regulatory elements from Arc mRNA in combination with synapse-targeting peptides^15^. Using an inducible Tet-On system for controlled expression, this approach enables the expression of tags, such as Venus and Hemagglutinin (HA), specifically at potentiated spines. Neurons were initially transfected with constructs (i) TreP-Arc3’-Nend-PSDTag-Venus-HA-Arc5’UTR; (ii) CAG-rtTA-IRES-TdTomato collectively referred to as SA-tdTomato; and (iii) pAAV.CAG.hChR2(H134R)-mCherry.WPRE.SV40 (ChR2-mCherry). Live imaging experiments using spinning disk (SD) microscopy show dendritic spines of transfected neurons A and B before illumination (Figures 3B). Spines that initially lack Venus fluorescence, progressively accumulate the fluorescent signal 60 and 90 min after optical stimulation, indicating synaptic strengthening. We also observed spines that emerged exclusively after illumination and exhibited robust Venus fluorescence 90 min later (Figure 3C). Overall, our finding indicates that both pre-existing and newly formed spines are strengthened in direct response to optical stimulation.

**Figure 3.**
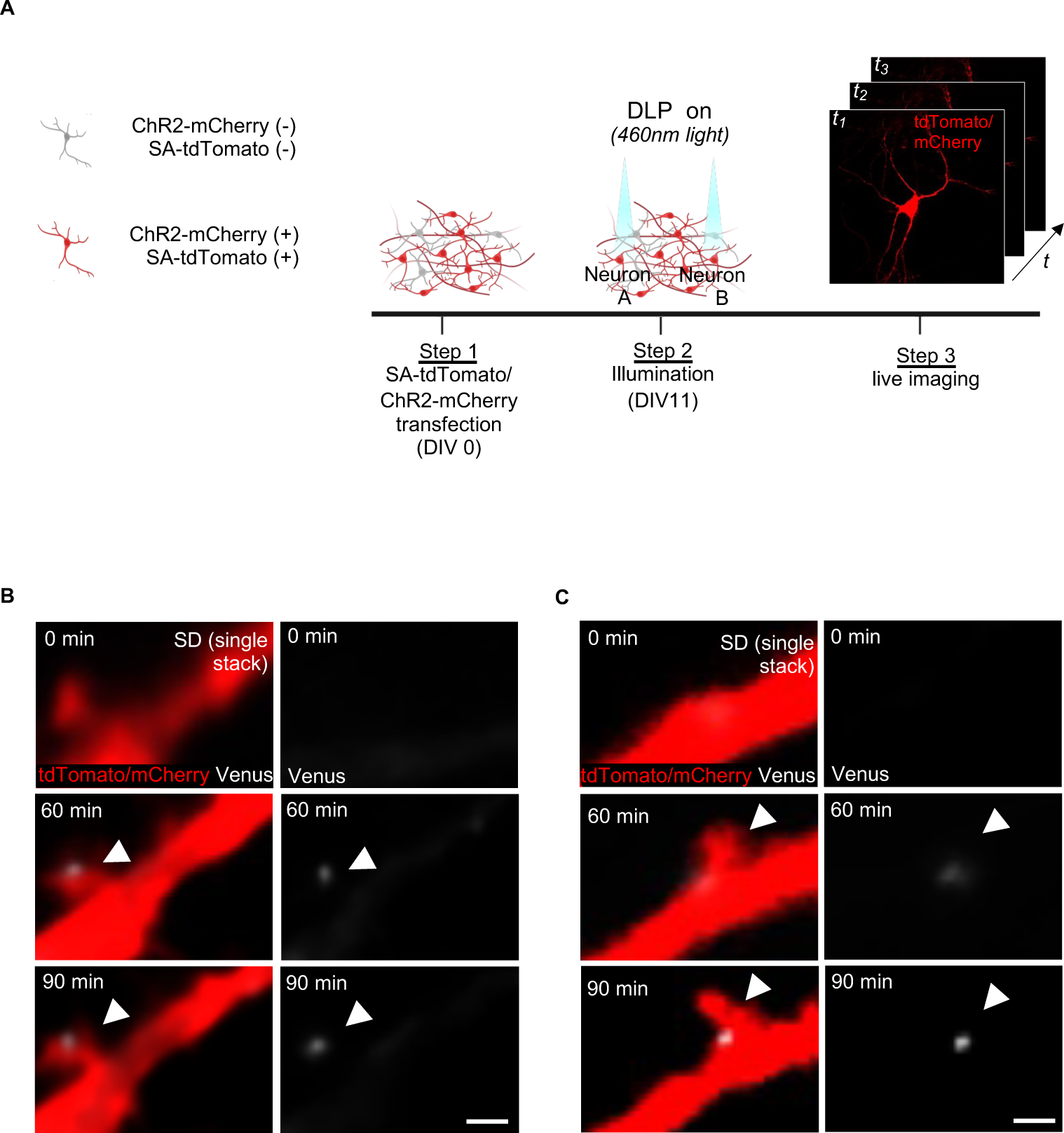
Tracking Venus tag accumulation in spine using live imaging. (A) Schematic diagram of the experimental procedures. (B) Dendritic spines expressing tdTomato gradually accumulate the Venus tag signal 60 and 90 minutes after the light stimulus. Arrowhead indicates HA+ spine. Scale bar: 1 μm. (C) Dendritic spines expressing tdTomato appeared and exhibited Venus tag fluorescence 60 and 90 minutes after illumination. Arrowhead indicates HA+ spine. Scale bar: 1 μm.

Quantitative analysis was performed by detecting HA immunofluorescence in neurons transfected with SA-tdTomato and transduced with AAV-ChR2-YFP 90 min after illumination (Figure 4A). The HA signal was analyzed using laser scanning confocal (LSC) microscopy (Figures 4C and S4) as well as structured illumination microscopy (SIM) (Figure 4D)^18–20^. Illuminated neurons A and B exhibited a significant increase in HA+ spines along their dendritic branches compared to non-stimulated neurons within and outside the FOV (Figure 4B). In contrast, neurons that were illuminated individually showed no similar increase in HA+ spine (Figures 4E and 4F). This indicates that the strengthening of dendritic spines relies on the coordinated activity of the prototypical two neurons, A and B, simultaneously stimulated within the network.

**Figure 4.**
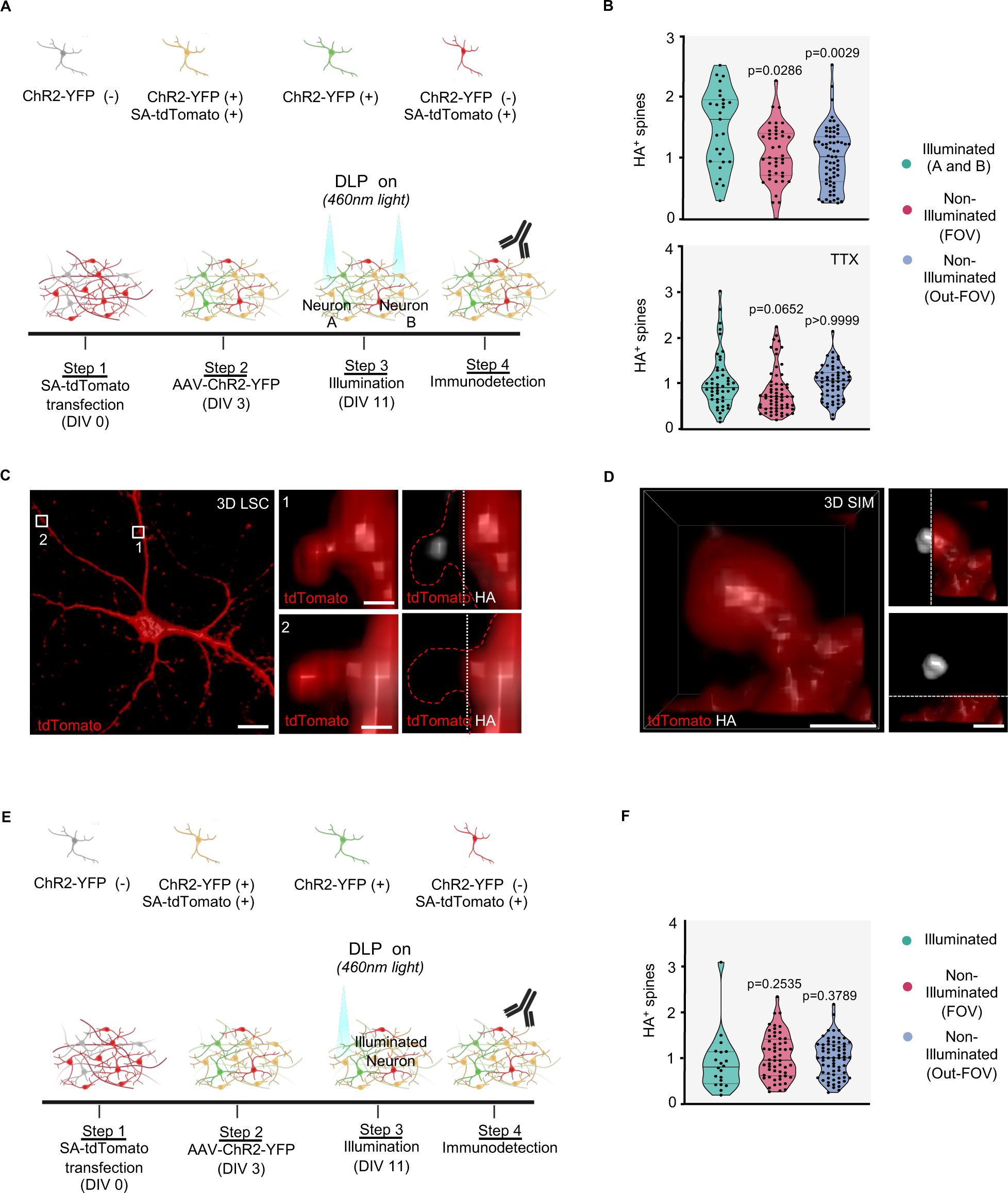
Quantitative analysis of HA tag accumulation in spines. (A) Experimental procedures for two-neuron illumination. (B) Upper, quantification of HA+ spines (%) in illuminated neurons A and B, non-illuminated neurons in the FOV and Out-FOV (n=27 dendrites for illuminated (A and B); n=42 dendrites for non-illuminated (FOV); n=67 dendrites for non-illuminated (Out-FOV); Kruskal-Wallis test). Lower, violin plot as in upper panel in the presence of TTX (n=40 dendrites for illuminated (A and B); n=61 dendrites for non-illuminated (FOV); n=58 dendrites for non-illuminated (Out-FOV); Kruskal-Wallis test). Data are normalized over the Out-FOV. (C) Left, 3D LSC images of a stimulated neuron expressing tdTomato. Scale bar: 10 μm. Right, the magnification of HA+ (1) and HA-(2) spines is shown; tdTomato signal was partially removed to unmask the HA signal. Scale bars: 1 μm. (D) Left, 3D SIM images of an HA+ spine. Right, the same spine is shown with the tdTomato signal partially (upper) or fully (lower) removed. Scale bars: 1 μm. (E) Experimental procedures for single-neuron illumination (F) Quantification of HA+ spines (%) in single illuminated neurons, non-illuminated neurons within the FOV and Out-FOV (n=27 dendrites for illuminated; n=42 dendrites for non-illuminated (FOV); n=67 dendrites for non-illuminated (Out-FOV); Kruskal-Wallis test). Data are normalized over the Out-FOV.

To validate this conclusion, we conducted experiments using light stimulation in the presence of Tetrodotoxin (TTX), which specifically blocks voltage-gated sodium channels and prevents action potentials in neurons, while leaving ChR2 activity unaffected^21^. The previously observed increase in HA+ spines of light-stimulated neurons A and B was abolished by TTX (Figure 4B), reinforcing the idea that synaptic strengthening occurs when target neurons receive synchronized pre- and post-synaptic inputs. This finding aligns with Hebb’s postulate on engram circuits.

Additional control procedures related to our experimental methods are reported in Figure S5 and Document S2.

### Patterned distribution of potentiated synapses

Dendritic spines are dynamic structures that modify their shape and strength in response to neuronal activity, shaping the wiring of neural circuits. We, therefore, investigated whether optical stimulation of the two-neuron modules can direct synaptic strengthening toward a specific pattern. To address this issue, we evaluated the spatial distribution of HA+ spines in illuminated neurons, analyzing their relative positioning within dendritic processes. We defined the orientation between neurons A and B as 0 degrees (0°) and evaluated the orientation of each dendritic process relative to this reference angle (Figure 5A). Dendrites showing an angle range of 0°-90° and 270°-360° were classified as IN-dendrites, while dendrites oriented in the opposite direction (angle range 90°-180° and 180°-270°) were classified as OUT-dendrites (Figure 5A). We then plotted the HA+ spines in each dendrite according to their IN-or-OUT categorization. Our data show a preferential distribution of HA+ spines toward IN-dendrites (Figures 5B and 5C). This polarized distribution was absent when neurons were illuminated with a weaker stimulation protocol (Figure 5C), which failed to induce synaptic potentiation (Figure S5B), or when exposed to TTX (Figure 5C). This suggests that the preferred orientation of HA+ spines is linked to the synchronized, reciprocal activity triggered by optical stimulation, and, thus, shapes their wiring. To support this conclusion, we demonstrated that the overall distribution of post-synaptic sites, marked by PSD95/tdTomato colocalization, did not significantly change between the dendrites of illuminated neurons (Figure 5D), suggesting that the observed changes in HA+ spines involve functional rather than anatomical reorganization.

**Figure 5.**
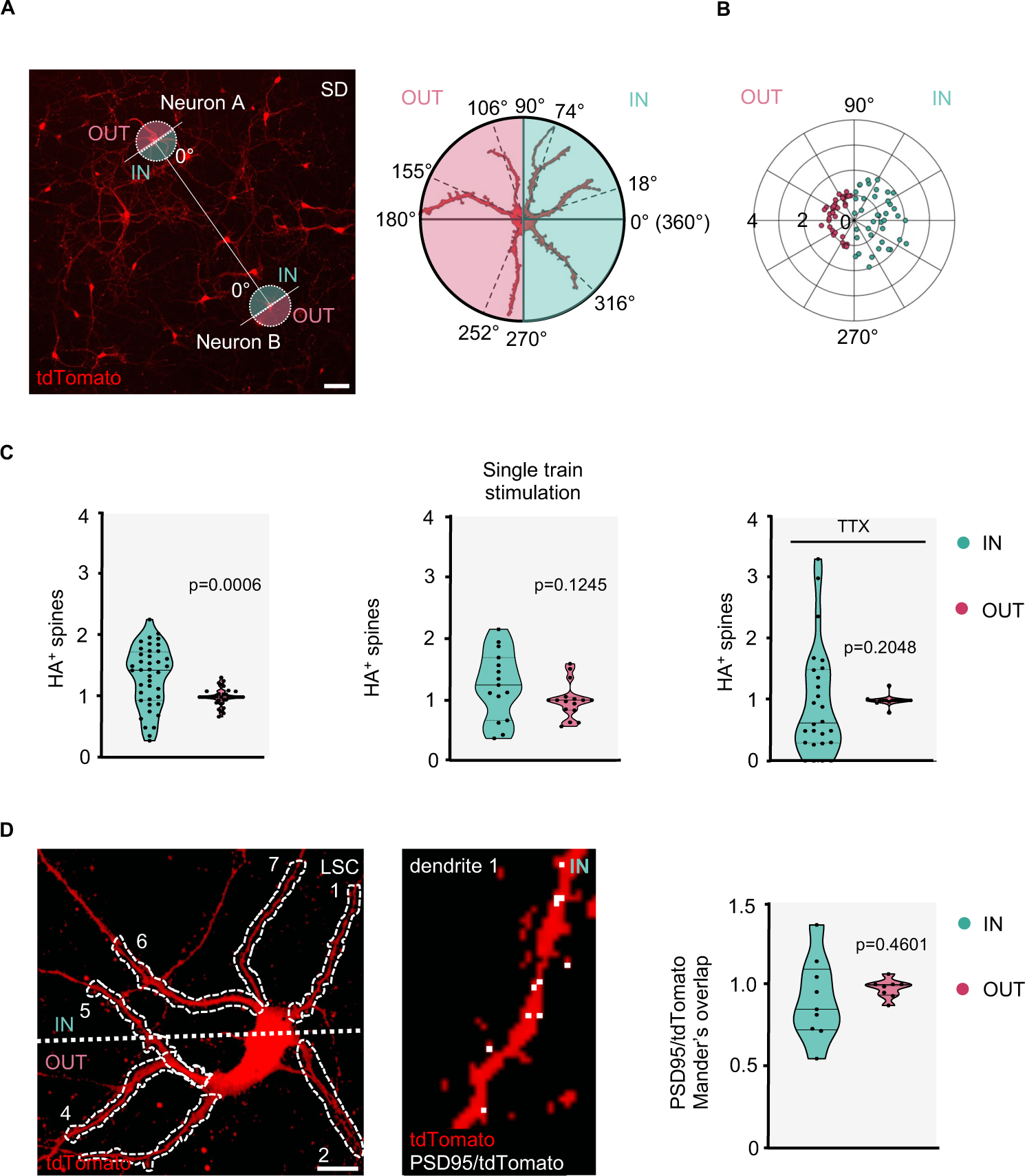
Spines distribution in light-stimulated neurons. (A) Left, SD image of tdTomato-expressing neurons within a network. The reference orientation between neurons A and B, is marked as 0 degrees (0°) on the polar plots. Dendrites falling within an angle range of 0° - 90° and 270° - 360° were classified as IN-dendrites (green), while dendrites within an angle range 90° - 180° and 180° - 270° are classified as OUT-dendrites (pink). Right, schematic representation of the polar plot displaying dendrites according to their oriented angles. Scale bar: 50 μm. (B) Polar plot illustrates the distribution of HA+ spines along the IN/OUT dendrites. (C) Left, HA+ spines quantification (%) of IN/OUT dendrites (n=40 dendrites IN; n=37 dendrites OUT; Mann-Whitney test). Middle HA+ spines quantification (%) of IN/OUT dendrites in neurons stimulated with 1 train stimulation (n=9 dendrites IN; n=8 dendrites OUT; Unpaired t-test). Right, HA+ spines quantification (%) of IN/OUT dendrites in the presence of TTX (n=25 dendrites IN; n=37 dendrites OUT; Mann-Whitney test). Data are normalized to the average of the “OUT” dendrites. (D) Left, LSC image of a tdTomato expressing neuron in which dendrites 1 to 7 (white dashed) are indicated. Two-time magnification of dendrite 1 showing PSD95/tdTomato co-localization. Right, quantification of PSD95/tdTomato colocalization in IN/OUT processes using Mander’s overlap coefficient. (n=9 dendrites for IN; n=8 dendrites for OUT; Unpaired t-test). Data are normalized to the average of the “OUT” dendrites. Scale bar: 10 μm.

Overall, our data capture the essence of Hebbian plasticity, often summarized by the axiom “neurons that fire together wire together”^22^ and provide robust conceptual support for our methodology.

### Direct targeting of synaptic ensembles

Our system allows precise scaling of DLP illumination to dendritic spine resolution (Figure S1C). Using this approach, we specifically illuminated spines on target dendrites (Figure 6A). In this process, the presynaptic afferents connected to the spines are also illuminated and optically stimulated, as they are engineered to express ChR2. This methodology allows the direct strengthening of synaptic contacts while minimizing overall neuronal stimulation.

**Figure 6.**
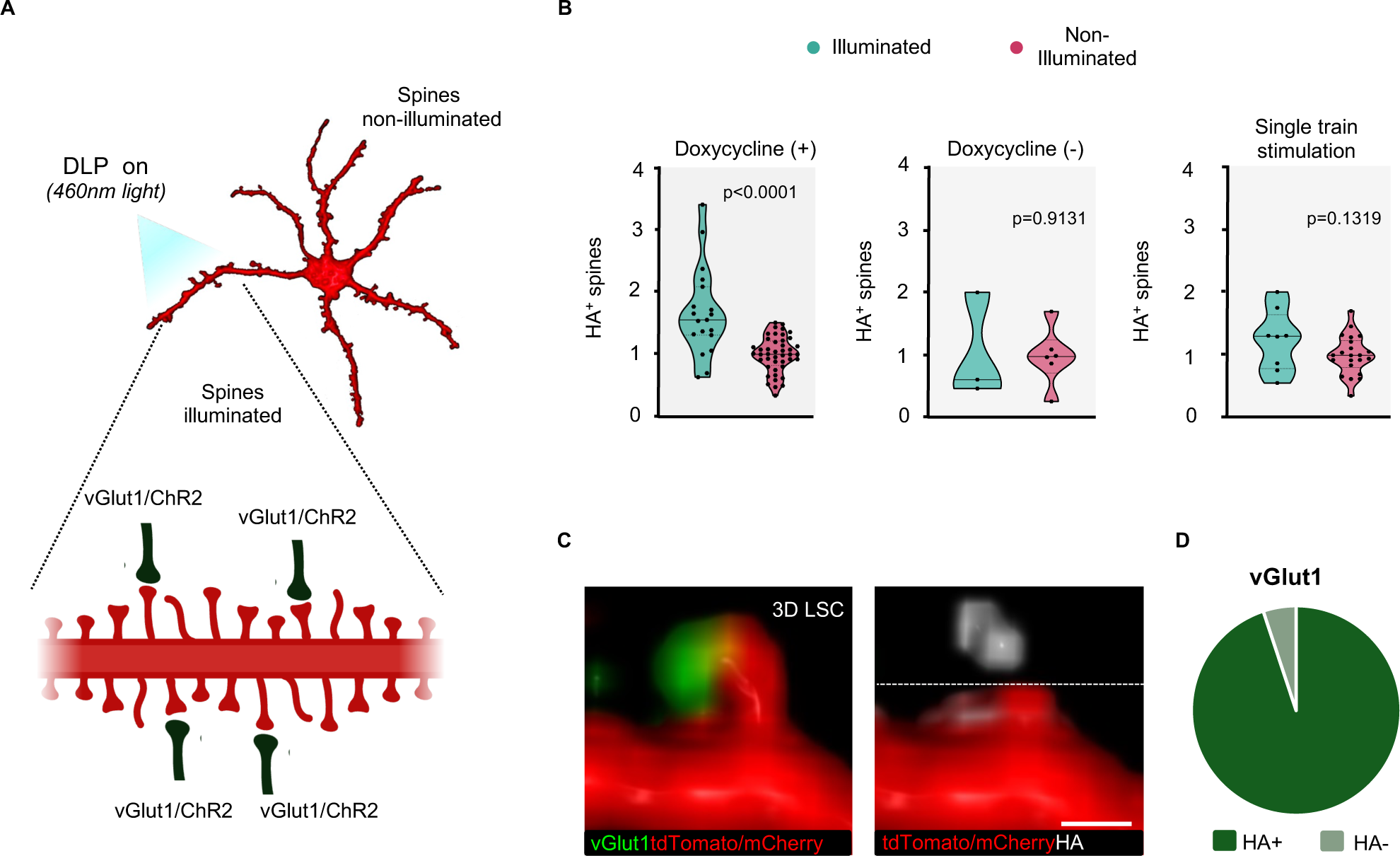
Induction of targeted synaptic strengthening. (A) Cartoon depicts the local illumination of a dendritic segment. Pre-synaptic afferents and post-synaptic structures expressing vGlut1/ChR2 are shown. (B) Left, quantification of HA+ spines (%) of illuminated dendrites in the presence (n=19 dendrites for illuminated; n=44 dendrites for non-illuminated dendrites; Unpaired t-test) and absence (n=3 dendrites for illuminated; n=3 dendrites for non-illuminated dendrites; Unpaired t-test) of doxycycline. Right, quantification of HA+ spines (%) of dendrites illuminated with single train stimulation and non-illuminated dendrites (n=8 dendrites for 1 single-light stimulation; n=22 dendrites for non-illuminated dendrites, Unpaired t-test). Data are normalized to the average of non-illuminated dendrites for each neuron. (C) Left, 3D LSC images of a spine expressing tdTomato/mCherry, positioned opposite to the vGlut1 signal. Right, the same spine with the HA signal unmasked. Scale bars: 1 μm. (D) Quantification of HA+ and HA-spines (%) positioned opposite to the vGlut1 signal.

Using a 60 X water immersion objective, we locally illuminate a portion of a dendrite in neurons expressing SA-tdTomato and ChR2-mCherry (Figure 6A). Following illumination, we observed a significant increase in HA+ spines in the illuminated regions after a latency of 90 min, compared to similar portions in the non-illuminated dendrites of the same neuron. Most of the spines (94.90%) showing proper pre-synaptic innervation, as indicated by vGlut1 immunolabeling, were also HA+ (Figure 6C and 6D). The increase in HA+ spines was not detected when Doxycycline was omitted or when a weaker stimulation protocol was applied (Figure 6B), as both conditions failed to promote synaptic potentiation (Figure S5B and S5C).

Overall, this experimental approach establishes a highly controlled environment for inducing synaptic strengthening in defined neuronal ensembles, allowing for precise manipulation and observation of the functional assembly and connectivity within neural networks. By targeting specific spines or clusters, we can systematically explore how various patterns of activity affect network assembly and the overall wiring of circuits, enabling the study of artificial synaptic engrams.

## Discussion

Our hybrid DLP system is a reliable tool for investigating the principles of synaptic ensemble strengthening in small-scale neuronal assemblies. It enables precise tracking and induction of artificial synaptic strengthening by integrating cutting-edge techniques such as optogenetics and synaptic imaging. This methodology is deeply anchored in Hebb’s foundational postulate on memory engrams, offering a solid theoretical framework that guides our research approach. At the core of our investigation is the reinforcement of synaptic strength between what we have termed ‘Hebbian’ neurons A and B. This critical process is essential for creating functional connectivity supporting the formation of neuronal assemblies within the brain^8,23^. The inherent simplicity of the two-neuron module, used as a building block for more complex circuits, offers an opportunity to explore the basic principles of engram circuits in situations where *in vivo* studies are not feasible.

Observational studies have indicated that changes in IEGs expression triggered by learning are closely associated with engram encoding and can predict future retrieval activities^17^. Common markers for identifying engram cells in living organisms include increased intracellular Ca^2+^ levels and the expression of cFos and Arc. Although these markers cannot define an engram *in vitro* - since an engram is identified through memory assessment *in vivo* - their presence in our system supports the identification of functional neuron ensembles that exhibit key characteristics of engram neurons. Likewise, the observed potentiation of spines between illuminated neurons A and B, as demonstrated through SA labeling, strongly validates our methodological approach. These findings not only confirm the system’s effectiveness in inducing, detecting, and quantifying synaptic changes between active neurons but also underscore its reliability in studying their spatial organization in synaptic ensembles.

The architectural organization of active synapses within an engram circuit is critical for understanding the wiring of engram neurons. Our study highlights that the specific orientations in which synaptic connections are strengthened play a crucial role in locating regions of heightened activity. Neurons subjected to patterned illumination formed connections that were not only strengthened but also directionally organized. This suggests an adaptive network mechanism that improves information processing efficiency. This supports the principle that “neurons that fire together wire together” emphasizing how synchronized neural activity contributes to memory engram formation by potentially integrating neurons across different brain areas more effectively. Our system proves to be a valid methodology for investigating the underlying principles of temporal and spatial organization within cell ensembles, offering deeper insights into the mechanisms of engram formation and function.

Moreover, the hybrid DLP system enables the precise illumination of individual pre- and post-synaptic sites, allowing for the direct formation of controlled synaptic ensembles. By shaping artificial synaptic ensembles, our system provides a powerful methodology for exploring principles underlying the architectures and configurations of synaptic engrams. This capability may bridge the gap between artificial intelligence models and biological research, offering a platform to test and validate theoretical predictions about neural connectivity. As such, it serves as a key resource for advancing our understanding of both natural and artificial neural networks.

After encoding, memory consolidation involves significant changes in the physical and chemical structure of engrams, including reinforced synaptic connections and altered gene expression^5,24^. Our methodology is well-suited for investigating these consolidation mechanisms. It allows precise control over stimulation patterns and durations to enhance synaptic plasticity, providing insights into the temporal dynamics of consolidation processing in neural circuits. Additionally, with advanced optical tools for protein manipulation, the system enables exploration of the spatial-temporal aspects of synaptic strengthening, crucial for understanding the dynamics of engram circuits and their key synaptic interactions.

In conclusion, the DLP system significantly advances our understanding of engram circuits by providing critical insights into the processes of synaptic encoding/consolidation and the temporal dynamics that underlie them.

## Acknowledgments

We thank Antonino Cattaneo for providing -TreP-Arc3’-Nend-PSDTag-Venus-HA-Arc5’UTR and CAG-rtTA-IRES-TdTomato constructs. We thank the MOF facility of the University of Trento. We thank Mattia Mancinelli for his help in setting up the DLP system and Ca^2+^ imaging analysis.

This project has received funding from the European Research Council (ERC) under the European Union’s Horizon 2020 research and innovation program, grant number 788793-BACKUP to L.P., Progetti di Ricerca di Rilevanza Nazionale (PRIN)-Bando 2017, grant number 2017HPTFFCPRIN to M.C, Fondazione Caritro to C.Z and European Union’s Horizon 2020 research and innovation program under the Marie Sklodowska-Curie grant agreement N. 101033260 (project ISLAND) to I.A.

## Author Contributions

M.C., B.V. and L.P. conceived the study; C.Z., A.M. and B.V. performed light-stimulation and Ca^2+^ imaging experiments; A.M and Y.H. prepared primary cultures; A.M. and B.V. performed immunohistochemistry; A.M., C.Z. and B.V. performed LSC, SD and SIM acquisitions; C.Z. implemented the DLP-SD setup; C.Z. implemented the code and performed Ca^2+^ analysis; C.Z. and A.M. performed spine analysis; I.A. and Y.H. performed MEA experiments and analyzed the data; M.C. and B.V. wrote the manuscript with input from C.Z., A.M. and L.P. and in collaboration with all other authors.

## Supplemental Information

Documents S1 and S2.

**Figures S1–S5.**

## Star Methods

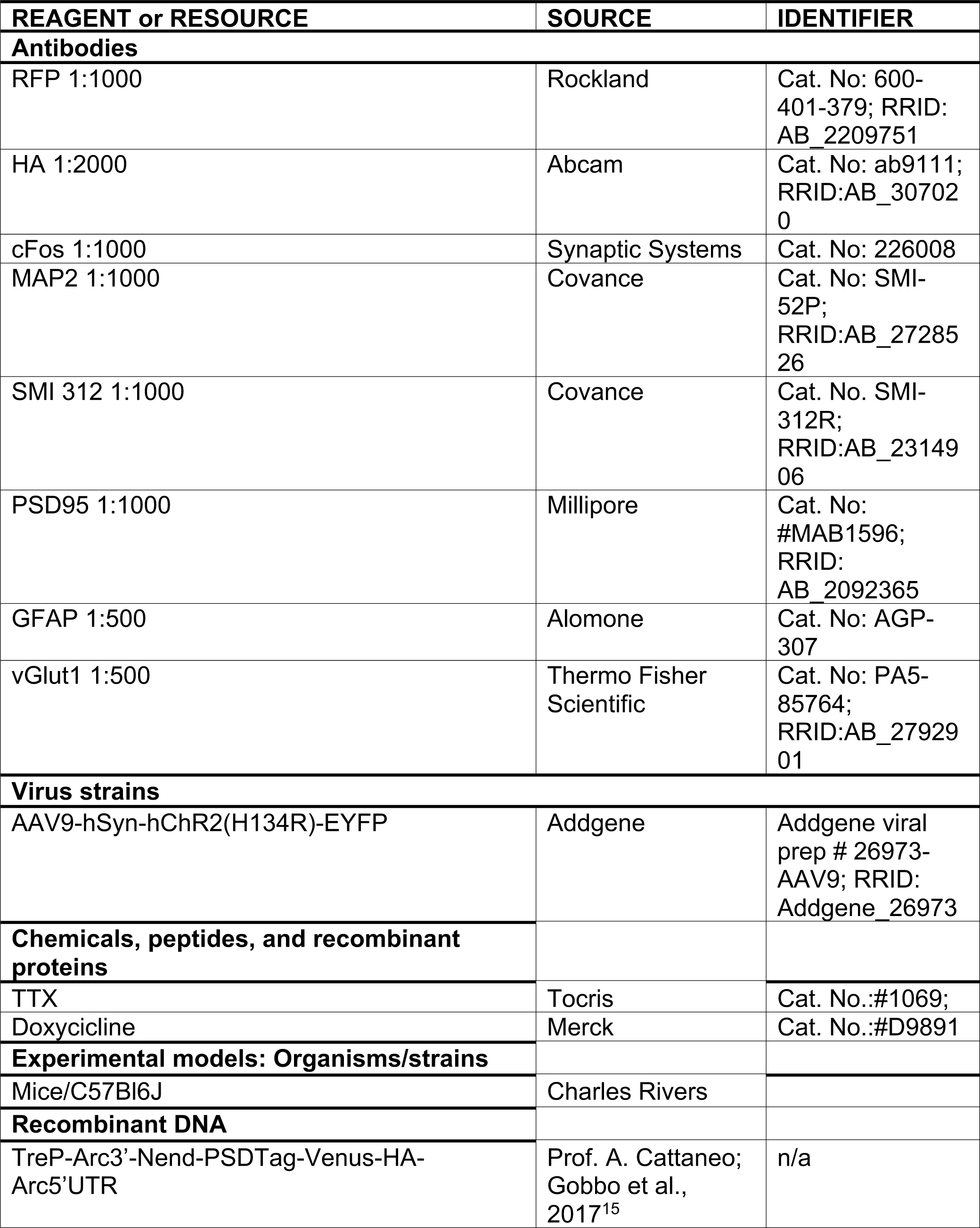
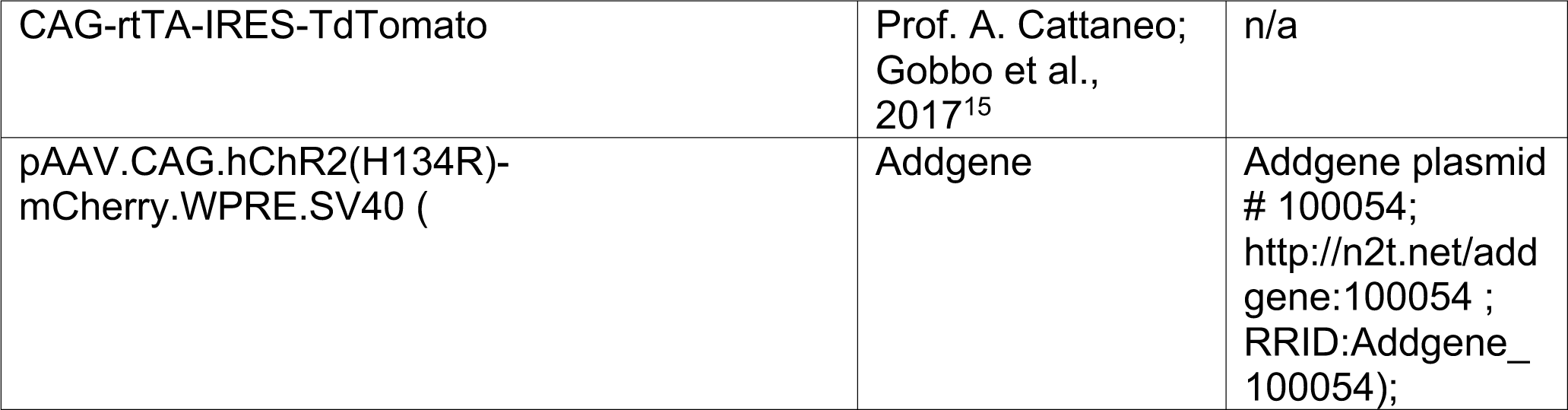

| REAGENT or RESOURCE | SOURCE | IDENTIFIER |
| --- | --- | --- |
| **Antibodies** |  |  |
| RFP 1:1000 | Rockland | Cat. No: 600-401-379; RRID: AB_2209751 |
| HA 1:2000 | Abcam | Cat. No: ab9111; RRID:AB_307020 |
| cFos 1:1000 | Synaptic Systems | Cat. No: 226008 |
| MAP2 1:1000 | Covance | Cat. No: SMI-52P; RRID:AB_2728526 |
| SMI 312 1:1000 | Covance | Cat. No. SMI-312R; RRID:AB_2314906 |
| PSD95 1:1000 | Millipore | Cat. No: #MAB1596; RRID: AB_2092365 |
| GFAP 1:500 | Alomone | Cat. No: AGP-307 |
| vGlut1 1:500 | Thermo Fisher Scientific | Cat. No: PA5-85764; RRID:AB_2792901 |
| **Virus strains** |  |  |
| AAV9-hSyn-hChR2(H134R)-EYFP | Addgene | Addgene viral prep # 26973-AAV9; RRID: Addgene_26973 |
| **Chemicals, peptides, and recombinant proteins** |  |  |
| TTX | Tocris | Cat. No.:#1069; |
| Doxycycline | Merck | Cat. No.:#D9891 |
| **Experimental models: Organisms/strains** |  |  |
| Mice/C57Bl6J | Charles Rivers |  |
| **Recombinant DNA** |  |  |
| TreP-Arc3'-Nend-PSDTag-Venus-HA-Arc5'UTR | Prof. A. Cattaneo; Gobbo et al., 2017<sup>15</sup> | n/a |
| CAG-rtTA-IRES-TdTomato | Prof. A. Cattaneo;<br>Gobbo et al.,<br>2017<sup>15</sup> | n/a |
| pAAV.CAG.hChR2(H134R)-<br>mCherry.WPRE.SV40 ( | Addgene | Addgene plasmid<br># 100054;<br>http://n2t.net/addgene:100054 ;<br>RRID:Addgene_100054); |

## Resource availability

### Lead contact

Further information and requests for resources and reagents should be directed to and will be fulfilled by the lead contact, Beatrice Vignoli.

## Materials Availability

This study did not generate new unique reagents.

## Experimental model

### Mice

All experiments were conducted in compliance with the Italian Animal Welfare legislation (D.L. 26/2014) that was implemented by the European Committee Council Directive (2010/63 EEC). The procedures were approved by local veterinary authorities and the Italian Ministry of Health (3FAF3.N.1JY). Mice were kept on a 12-hour light/dark cycle and had ad libitum access to food and water.

## Method details

### Primary cell culture

Cortices were isolated from C57BL6 mouse embryos at embryonic day 17 (E17). In brief, cortical tissues were collected and placed in ice-cold Hank’s balanced salt solution (HBSS) with 10 mM HEPES. The tissues were then incubated with 0.25% trypsin-EDTA at 37°C for 20 min. Subsequently, trypsin was carefully aspirated, and the tissues were washed twice with a phosphate-buffered saline (PBS). To neutralize the trypsin, 1 ml Dulbecco’s modified Eagle’s medium (DMEM), containing 10% fetal bovine serum (FBS) and penicillin-streptomycin 10000 U/ml (P/S) was added. The cortical cells were dissociated using a P1000 pipette, collected by centrifugation (1900 RPM for 5 min), and resuspended in DMEM supplemented with 10% FBS and P/S.

The cells were eventually transfected with 1.5 μg of each DNA construct using the Amaxa Nucleofector system, following the manufacturer’s guidelines. After transfection, the cells were plated on glass coverslips pre-coated with poly-L-lysine (0.1 mg/ml). Three hours after plating, the medium was replaced with Neurobasal supplemented with B-27, P/S, and Glutamax. Half of the culture medium was refreshed every 3-4 days. Cells transfected with SA-tdTomato were treated with 1μg/ml of doxycycline for 24 hours before light stimulation.

In some experiments, cells were infected at DIV3 with the adeno-associated virus AAV9-hSyn-hChR2(H134R)-EYFP (AAV-ChR2-YFP).

In some experiments, TTX (1μM) was applied 20 min before light stimulation.

### Constructs and viral vectors

The following constructs were used:

-TreP-Arc3’-Nend-PSDTag-Venus-HA-Arc5’UTR^15^;

CAG-rtTA-IRES-TdTomato^15^;

pAAV.CAG.hChR2(H134R)-mCherry.WPRE.SV40 was a gift from Karl Deisseroth (Addgene plasmid # 100054; http://n2t.net/addgene:100054; RRID:Addgene_100054);

The following viral vector was used:

-AAV9-hSyn-hChR2(H134R)-EYFP was a gift from Karl Deisseroth (Addgene viral prep # 26973-AAV9; http://n2t.net/addgene:26973; RRID: Addgene_26973).

### Experimental plan for DLP light stimulation experiments

At DIV11 cortical neurons were placed in a 35 mm dish with a dark bottom, filled with HBSS supplemented with 2 mM CaCl_2_ and 1 mM MgCl_2,_ for microscopy. Neurons were selected for individual excitations based on the expression of tdTomato (reporting SA) and YFP or mCherry (reporting ChR2). Preference was given to neurons with at least two processes not oriented in a specific direction. Light excitations were administered using a pattern of 10 trains of 13 light pulses delivered at 100 Hz, repeated at 0.5 Hz with an intensity of 11±1 mW/mm^2^. When two light spots were used, they were spaced 500-1200 μm apart. After light stimulation, coverslips were returned to the incubator with culture media for 90 min, then fixed with cold 4% paraformaldehyde (PFA) for 20 min. Non-illuminated cultures were kept in the dark throughout the entire experimental period.

### Immunocytochemistry

Fixed neurons were permeabilized with 0.1% Triton X-100 in PBS for 20 min and blocked with 3% bovine serum albumin (BSA) in PBS for 60 min. Neurons were incubated with primary antibodies diluted into 3% BSA at 4°C overnight. After incubation, the cells were washed several times with 3% BSA and then incubated with fluorophore-conjugated secondary antibodies for 2 h at room temperature. Following this, the cells were washed with PBS, counterstained with DAPI (Invitrogen), and mounted with Aqua Polymount (Polyscience Inc.).

### Spinning disk, confocal and structured illumination microscopies

The microscopy setup, purchased from Crisel Instruments Company, includes the X-Light V2 top-performance confocal spinning disk module and an integrated VCS (Video Confocal Super-resolution) module based on structured illumination microscopy (SIM). The SIM module achieves a lateral resolution of 115 nm and an axial resolution of approximately 250 nm. The system is configured in an upright orientation and incorporates 7 solid-state LED sources with a wavelength range from 395 nm to 640 nm, arranged in the Lumencore Spectra X LED-based light engine. The imaging system is equipped with a Prime Back-side Illuminated (BSI) Scientific CMOS camera with 2048×2048 pixels and a pixel area of 6.5μmx6.5μm. Three objectives are integrated into the system: a dry long-working-distance 10 X (LMPLFLN10XLWD, NA 0.25, WD 21mm), a water immersion 60 X (LUMPLFLN60XW, NA 1, WD 2mm), and an oil 100 X (UP objectives UPLSAPO100XO, NA 1.4, WD 0.13 mm). The confocal setup is equipped with spinning disks with double pinhole patterns: one with 40 μm holes and another with 70 μm holes. The 70 μm disk was used for all images in this study.

In SIM mode, the specimen is illuminated with a multipoint beam by filtering the excitation light through a mask and scanning in orthogonal dimensions. Image stacks were acquired with a format of 2048 X 2048 pixels, a z distance of 150 nm, and 36 raw images per plane (multiple acquisition mode x-y grid scan) using a 100 X oil objective. The 16-bit depth SIM raw data were computationally reconstructed with the Metamorph software package. Super-resolution algorithms are then employed to compute the final super-resolved image using NIS software.

### Digital light processor device

A Digital Light Processor device (DLP) E4500 purchased from E.K.B Technologies Ltd. equipped with 3 LEDs, optics, a WXGA DMD (Wide Extended Graphics Array Digital Micromirror Device), and a driver board was integrated into the EPI-fluorescence port of SD microscope. The DLP’s light engine can generate approximately 150 lumens at 15 W of LED power consumption. Specifically, a blue LED with a wavelength of 460 ± 14 nm and a power output of 600 mW was utilized. The DMD features 1039680 mirrors arranged in a diamond array configuration of 912 columns by 1140 rows, enabling patterned illumination using preloaded and custom patterns. The system supports temporal pulsing of light with internal and external triggers, with a minimum exposure time ranging from 235 μs to 8333 μs, depending on the bit depth of the projected image. It handles 1 to 8-bit images at a resolution of 912 columns × 1140 rows, ensuring pixel accuracy where each pixel corresponds to a micromirror on the DMD. The emitted light from the DLP system is aligned with the microscope’s optical path via the system’s rear port. A dichroic mirror (Chroma T505lpxr-UF1) serves as a high-pass filter, reflecting wavelengths smaller than 505 nm. This setup enables simultaneous DLP illumination and signal recording at longer wavelengths.

To determine the light power reaching the specimen, we employed a power meter beneath the microscope objective (Ophir Nova, PD300).

### Laser scanning confocal microscopy

Confocal imaging was performed using a laser-scanning motorized confocal system (Nikon A1) equipped with an Eclipse Ti-E inverted microscope and four laser lines (405, 488, 561 and 638 nm). Z-series images were taken with an inter-stack interval of 0.3 or 0.5 μm using a 60X oil immersion objective (Nikon APO 60x 1.4).

### Calcium imaging

At 10-13 DIV cells were treated with X-Rhod-1 AM (1μM) in HBSS supplemented with CaCl_2_ (2mM) and MgCl_2_ (1mM) at 37°C for 30 min. Subsequently, the cells were washed with supplemented HBSS at 37°C. After 30 min, coverslips were positioned under the spinning disk microscope in a 35 mm dark bottom dish with supplemented HBSS.

The FI of X-Rhod-1 was monitored by acquiring time-lapses at 1.5 Hz for about 3 min. At frame 100 of the timelapse, LTP light stimulation (10 trains of 13 light pulses delivered at 100 Hz, repeated at 0.5 Hz) from the DLP was applied. During DLP stimulation, the camera was triggered to acquire only during intervals when the DLP light was off (i.e., between the 10 light trains). The resulting timelapse file was then analyzed using a custom MATLAB program designed to identify neuronal bodies (ROIs) and extract calcium traces based on X-Rhod-1 fluorescence in these ROIs over time.

### Automated detection

For region of interest (ROI) identification, the process begins with creating an average image from all frames captured before excitation. This average image is divided into 50 intensity levels, used as minimum thresholds for generating binary images. To isolate neuronal cell bodies, the regionprops function in MATLAB is used to extract essential information from the analyzed regions. This information encompasses parameters such as area, centroid coordinates, axis dimensions, and intensity values; the process is performed on all the binary images and ROIs duplicates are then removed by evaluating the distances between their centroids.

### Ca^2+^ signal normalization

To process the multi-tiff images, we perform the following steps for each frame:

1. Calculate the average pixel intensity within identified ROIs to create temporal traces of calcium fluorescence.
2. To eliminate baseline fluorescence noise, we normalize the entire trace using a linear fit that extends up to the excitation point. This helps in setting a consistent baseline for comparison.
3. To distinguish calcium response traces from the noise, we use specific parameters: ΔF/F: represents the ratio of the difference (ΔF) between the maximum fluorescence level during light stimulation and the baseline fluorescence level (F), which is the average of all frames before stimulation, to F. Σ: indicates the maximum absolute variation of the fluorescence trace before the onset of light stimulus.
4. To classify traces as responses, we apply threshold values to these parameters. Specifically, we require: (i) ΔF/F to be greater than 0.03, indicating a significant increase in fluorescence during stimulation; (ii) 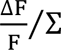 to be greater than 1.2, ensuring that the increase in fluorescence is significantly higher than any potential pre-stimulation fluctuations.

We classified a trace as a response when calcium indicator fluorescence changed above thresholds at the stimulation time. We calculated ΔF/F to quantify the change in fluorescence in these responsive neurons.

### Quantification of HA+ spines

Imaging processing was conducted using the NIS Elements software from Nikon. Confocal images underwent cropping, focusing on segments of dendrites proximal to the cell body within 30-60 μm range. Subsequently, spines along the dendritic processes were morphologically identified using the tdTomato cellular filler. Manual quantification of spines expressing the HA signal (HA+) was performed on single stack images, and the results were expressed as a percentage relative to the total number of spines for each cropped dendrite. Cells subjected to light stimulation were categorized as “illuminated” neurons while neighboring non-illuminated neurons within the field of view were designated as “non-illuminated” cells. Cells located far from the stimulated area were classified as “out-FOV”. The percentage of HA+ spines was normalized over the outFOV values.

To investigate the distribution of HA+ spines within the network, polar plots were generated to map the orientation of dendritic processes in stimulated cells and their corresponding percentage of HA+ spines. Specifically, for each stimulated cell, angles were measured between the imaginary line connecting the centers of cell bodies (defined as 0°) and the direction of the proximal part of the dendrites. Subsequently, polar plots were created associating each dendrite’s direction (angle) with its percentage of HA+ spines. The data were then categorized into “IN” and “OUT” processes based on their location in the first/fourth quadrant (facing the second neuron) of the polar plot or the second/third quadrant, respectively. For comparative analysis, for each cell the percentage of HA+ spines for all dendrites was normalized on the average of “OUT” processes.

### Live imaging experiments

Live imaging experiments were performed at DIV 13 on cultures transfected with SA and ChR2-mCherry. Coverslips were placed in a dark bottom 35 mm dish containing HBSS, CaCl_2_ (2mM) and MgCl_2_ (1mM). The dish was placed in the Okolab stage incubator system (H301-UP), equipped with H301-T-UNIT-BL-PLUS electric top stage incubation system, maintaining a constant temperature of 37°C. Cells expressing tdTomato/mCherry with clearly defined dendrites and visible spines were selected for observation. Two neurons were selected for spot LTP-like illumination using the DLP and a 10 X objective with an 11 ± 1 mW/mm^2^ intensity. The expression of Venus and tdTomato/mCherry was monitored using the 60 X water immersion objective, acquiring confocal images before, 60 min and 90 min after stimulation. Images were acquired by averaging 4 images for each z-stacks at 0.5 μm step size.

### Multi Electrode Array (MEA) experiments

A MEA-2100mini system from Multichannel Systems GmbH (MCS) was used to record the electrical activity of neuronal cultures. The cultures were plated on electrode array chips (60MEA-200/30iR-Ti-gr by MCS) featuring 60 titanium-nitride electrodes embedded in glass and enclosed by a glass ring. The electrodes have a diameter of 30 μm and a pitch of 200 μm. The chips are placed on a head-stage device that samples the culture’s electrical signals. The collected signals are amplified and filtered by a signal collecting unit (SCU), then digitalized and read by software.

In our experimental procedure, the DLP system delivered optical stimulation within a circular area with a diameter of approximately 200 μm. Initially, neurons were exposed to a baseline illumination pattern, featuring 20 ms light pulses at a frequency of 0.5 Hz for 10 minutes. Then, to induce long-term potentiation (LTP), neurons underwent a stimulation regimen that included 10 sets of 13 light pulses delivered at 100 Hz, with a repetition frequency of 0.5 Hz. After the LTP stimulus, neurons were subjected to baseline illumination patterns at four different time points: 0, 20, 40, and 60 min post-stimulation.

The recording is performed using MCS experimenter software, where the signals can be digitally filtered, inceptively analyzed, and tracked in real-time. The signals were sampled at 20KHz. The recorded files are saved and exported for subsequent offline analysis to identify neuronal spikes. Initially, raw signals underwent digital filtering with a band-pass Butterworth filter with cutoff frequencies set at 0.3 and 3 KHz. Spikes were detected using a Precise Timing Spike Detection (PSTD) algorithm^25^. To analyze the responses of the neuronal network to light stimulation, we used post-stimulus time histograms (PSTHs) at each electrode, counting the number of spikes within predefined time bins of fixed width and distance relative to the light pulse, averaged over the number of pulses. Electrodes were excluded from the analysis if they exhibited very low activity or unstable behavior, based on two criteria: an activity threshold and a stability threshold. The activity threshold was defined as an area under the PSTH curve of less than ‘1’ during the baseline step. To assess stability, the baseline recording was repeated without applying a tetanus stimulus in between, and the change in the PSTH area between the two recordings was calculated: an electrode was excluded from the analysis if the absolute change, |ΔP|/P (where P is the PSTH area of the reference baseline), exceeded 20%. For the selected electrodes, the efficacy of potentiation was calculated as the change in the PSTH area (ΔP/P) between the pre- and post-LTP pattern exposure phases. The electrodes indicating potentiated activity were defined as those that exceed the +20% stability threshold ^26^. The Potentiation Index (PI) was subsequently determined as the percentage of the channels that displayed an efficacy higher than the selected threshold at each time step among the selected channels, providing valuable insights into the progression of potentiation over time.

## Quantification and statistical analysis

All statistical analysis were performed using GraphPad Prism version 10.0.0, GraphPad Software, Boston, Massachusetts USA, www.graphpad.com PRISM. All violin plots depict median (continuous line) and quartiles (dashed line). All data were run for the Shapiro-Wilk normality test. Statistical comparisons were performed using a t-test or one-way ANOVA test for normally distributed data and Mann-Whitney or Kruskal-Wallis tests for non-normally distributed data. Details on the sample number n, the type of statistical test used, and the significance of the data are provided in the figure legends. A p-value <0.05 was considered statistically significant.

