## Supplemental Text and Figures for "Mapping synaptic ensembles through *in vitro* functional cell assemblies"

**Document S1. DLP- based system**

*Light Delivery.* Our DLP-based system achieves varying levels of resolution, defined as the full-width half maximum (FWHM) of the illuminated region on the sample plane when a single pixel of the DLP matrix is activated. Specifically, with a 10 X air objective, the resolution is 5.4 ± 0.2 μm when the aperture stop (AS) is fully closed to minimize optical aberrations (Figure S1C). With the AS open, the resolution is slightly reduced to 8 ± 5 μm. Using a 60 X water immersion objective, significantly improves the resolution, reaching 0.90 ± 0.07 μm with the AS closed (Figure S1C) and 1.1 ± 0.6 μm with the AS open. Based on this resolution, the 10 X objective is ideal for illuminating cell bodies, while the 60 X objective is suited for illuminating dendritic spines.

The light distribution on the sample plane was initially not uniform (Figure S1A), causing variations in light intensity experienced by neurons at different positions within the FOV. To correct this issue, we rectified the non-uniform light distribution mapping the widefield light intensity distribution on the sample plane and we used it as a reference for normalizing the desired black-and-white light pattern. This pattern was then loaded into the DLP software as an 8-bit image. Additionally, we verified that configuring the DLP with a high bit depth did not affect the temporal patterns used in our experiments. This approach allowed us to enhance the uniformity of the light distribution, especially at the center of the FOV (Figure S1A). To measure the light power reaching the specimen, we used a power meter placed beneath the microscope objectives. The intensity at the sample plane was calculated by adjusting for the illumination area. We ensured a consistent average intensity of 11 ± 1 mW/mm² across the entire FOV. Careful adjustment of the aperture (AS) and field stop (FS) positions confined the illumination to the desired FOV, improving the correction of optical aberrations and contrast.

*Pattern Characterization.* To achieve single-cell excitation within the neuronal network, we designed localized light spots at specific positions for each experiment. These spots were set to have diameters ranging from 10 to 40 µm (Figure S1B). Since the average distance between ChR2-expressing neurons in our cultures is approximately 170 µm, the dimensions of a single spot, centered on a cell body are optimal for precise single-cell excitation. Regarding contrast, defined as (max-min)/min*100, the average contrast of the single spot across the FOV was (5 ± 2) x 10^4 relative to the background signal. However, when multiple spots were projected simultaneously (24 spots; Figure S1D), the background signal increased, resulting in a contrast value of (16 ± 5) x 10^3.

*Two-spots illumination.* Most of our study focused on two-spot illumination using a 10 X objective. To determine the minimum distance at which two spots are considered separated, we projected pairs of spots at different distances and analyzed their intensity profiles. Specifically, we evaluated the ratio between the valley intensity and the average of the two peak intensities (valley/peak). We accounted for potential aberrations that could affect the minimum distance between spots differently in the vertical (Figure S1E) and horizontal (Figure S1F) directions. To address this, we averaged the valley/peak values in both the vertical and horizontal directions, finding that the separation reached the value of 0 at approximately 120 µm (Figure S1G).

*Light stimulation of dendritic spines.* The illuminated area had a diameter of 45 ± 8 μm and was positioned 18 ± 4 μm away from the cell body.

*Light to Noise Characterization*. To eliminate potential interference or noise from multiple reflections between the glass coverslip, where neurons are plated, and the Petri dish containing the buffer solution, we performed all characterizations on a reflective surface placed at the sample plane and immersed in distilled water. Although cell bodies can influence light transmission, and the coverslip can cause multiple reflections between the glass and the dish's plastic, we considered this noise negligible compared to the direct light emitted by the DLP. Nonetheless, to further minimize reflections at the dish-air interface during experiments, we used dishes with black bottoms. This approach helped to reduce potential light noise, ensuring the accuracy of our observations.

**Document S2. Additional control experiments**

To further strengthen the reliability of our experimental approach, we conducted additional control experiments. These involved broad, wide-field illumination that encompassed the entire network within the FOV (Figure S5A). Under these conditions, we performed the following experiments:

*Single train*. To verify that the increase in HA+ spines in our system relies on a specific light pattern stimulation, we performed experiments using a single train of 13 light pulses instead of the full stimulation protocol necessary to induce LTP. With this weaker stimulation protocol, the illuminated neurons displayed a similar number of HA+ spines as the non-illuminated neurons (Figure S5B).

*Doxycycline removal*. The SA technology is based on an inducible Tet-ON system, where HA genes are controlled by a promoter (TREp) that responds to doxycycline. When doxycycline is present, it binds to the reverse tetracycline-controlled transactivator (rtTA), which is under the control of the human synapsin promoter (hSYNp). This binding causes a structural change in rtTA, activating the transcription of HA gene. To confirm the system's selectivity, neurons were exposed to light stimulation with or without doxycycline. The illumination pattern and intensity (11 ± 1 $mW/mm^2$) were consistent with those used for LTP induction. In the presence of doxycycline, neurons within the illuminated field showed an increase in HA+ spines compared to non-illuminated neurons. In contrast, there was no increase in HA+ spines in optically stimulated neurons without doxycycline (Figure S5C).

*Procedural stages*. In our experimental protocol, cells undergo several distinct stages. First, they are placed in an incubator and then transferred to a microscope to visualize neurons transduced with ChR2-YFP. Once a FOV is selected, neurons are exposed to illumination by a DLP. To eliminate potential interference from light sources other than the DLP on the activation of ChR2-YFP-expressing neurons, we performed HA immunodetection on neurons in the incubator (condition 1), before DLP illumination (condition 2), and after DLP illumination (condition 3) (Figure S5D). Our analysis shows a significant increase in HA+ spines only when neurons are subjected to field illumination from DLP.

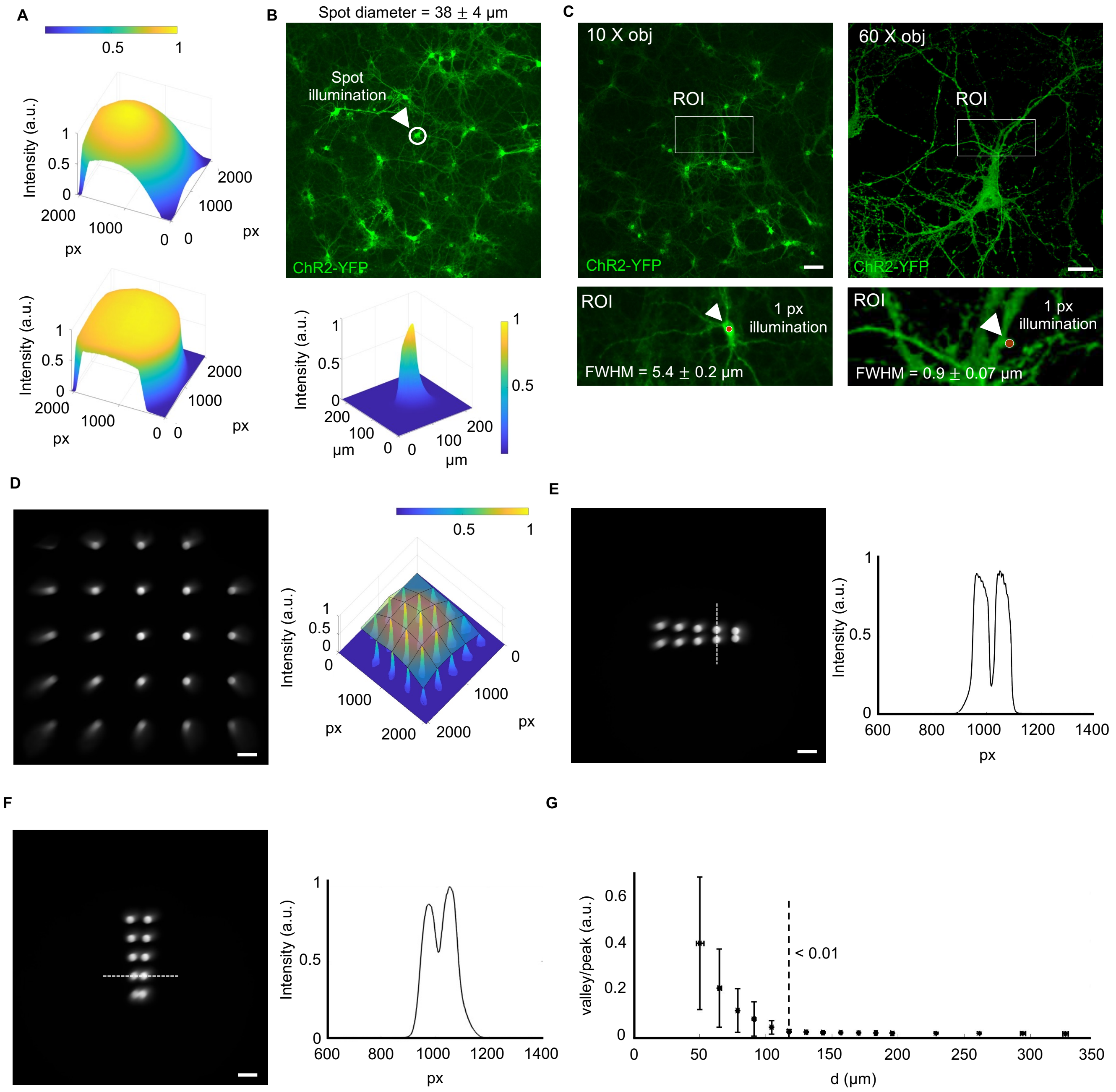
A
B
Spot diameter = 38 ± 4 µm
Spot illumination
ChR2-YFP
C
10 X obj
60 X obj
ROI
ChR2-YFP
ROI
1 px illumination
FWHM = 5.4 ± 0.2 µm
FWHM = 0.9 ± 0.07 µm
Intensity (a.u.)
px
µm
D
E
F
G
valley/peak (a.u.)
d (µm)
< 0.01

**Figure S1. DLP-based system. Related to Figure 1 and Document S1**

(**A**) Light intensity distribution on the sample plane before (upper) and after (lower) correction.

(**B**) Upper, ChR2-YFP-expressing neurons in the FOV. The ROI (circle) indicates a typical spot used for neuron illumination in the DLP experiments. Lower, light intensity distribution of the typical spot illumination. Scale bar: 50 μm.

(**C**) Upper, ChR2-YFP-expressing neurons in the FOV using a 10 X objective (left; scale bar: 50 μm) and 60 X objective (right; scale bar: 10 μm). Lower, ROIs shown at three times magnification, depicting 1-pixel illumination using 10 X objective (left) and 60 X objective (right). Arrowheads indicate the illuminated areas.

(**D**) Multiple spots of illumination in the FOV (left), with the corresponding intensity distribution (right). Scale bar: 100 μm.

(**E**) Left, a series of two illuminated spots projected at varying vertical distances from each other. Right, light intensity profile measured across the two spots, as indicated by the dashed line in the left panel. Scale bar: 100 μm.

(**F**) Left, a series of two illuminated spots projected at varying horizontal distances from each other. Right, light intensity profile measured across the two spots, as indicated by the dashed line in the left panel. Scale bar: 100 μm.

(**G**) Valley/peak ratio values averaged in the vertical and horizontal directions plotted against distances between spots. The valley/peak ratio values from the dashed line are less than 0.01.

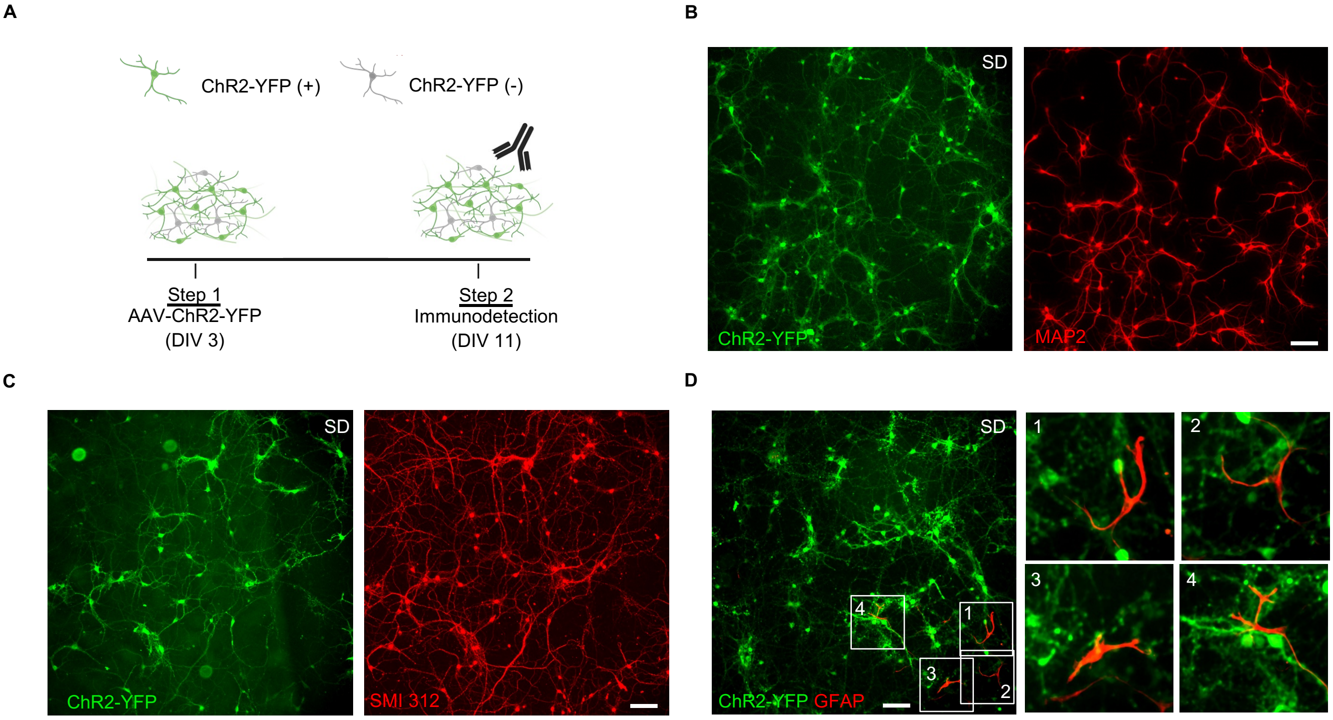
A
ChR2-YFP (+)
ChR2-YFP (-)
Step 1
AAV-ChR2-YFP
(DIV 3)
Step 2
Immunodetection
(DIV 11)
B
SD
ChR2-YFP
MAP2
C
SD
ChR2-YFP
SMI 312
D
SD
ChR2-YFP GFAP
1
2
3
4

**Figure S2. ChR2 expression and marker-characterization in cortical neurons. Related to Figure 2.**

(**A**) Schematic diagram of the experimental procedures.

(**B**) Representative SD images of cortical neurons expressing ChR2-YFP (left) and Microtubule-Associated Protein 2 (MAP2; right). The images depict a neuronal network consisting of independent cell bodies that are homogeneously distributed within a typical FOV, showing multiple dendrites with intricate branching processes.

(**C**) Representative SD images of cortical neurons expressing ChR2-YFP (left) and Purified anti-Neurofilament Marker (SMI 312; right). The SMI 312 staining reveals a complex axonal network connecting most neurons within the FOV.

(**D**) Representative SD images of cortical neurons expressing ChR2-YFP and Glial Fibrillary Acidic Protein (GFAP). ROIs 1 to 4 show a three-time magnification of GFAP+/ChR2-YFP- cells. Scale bars: 50 $\mu$m.

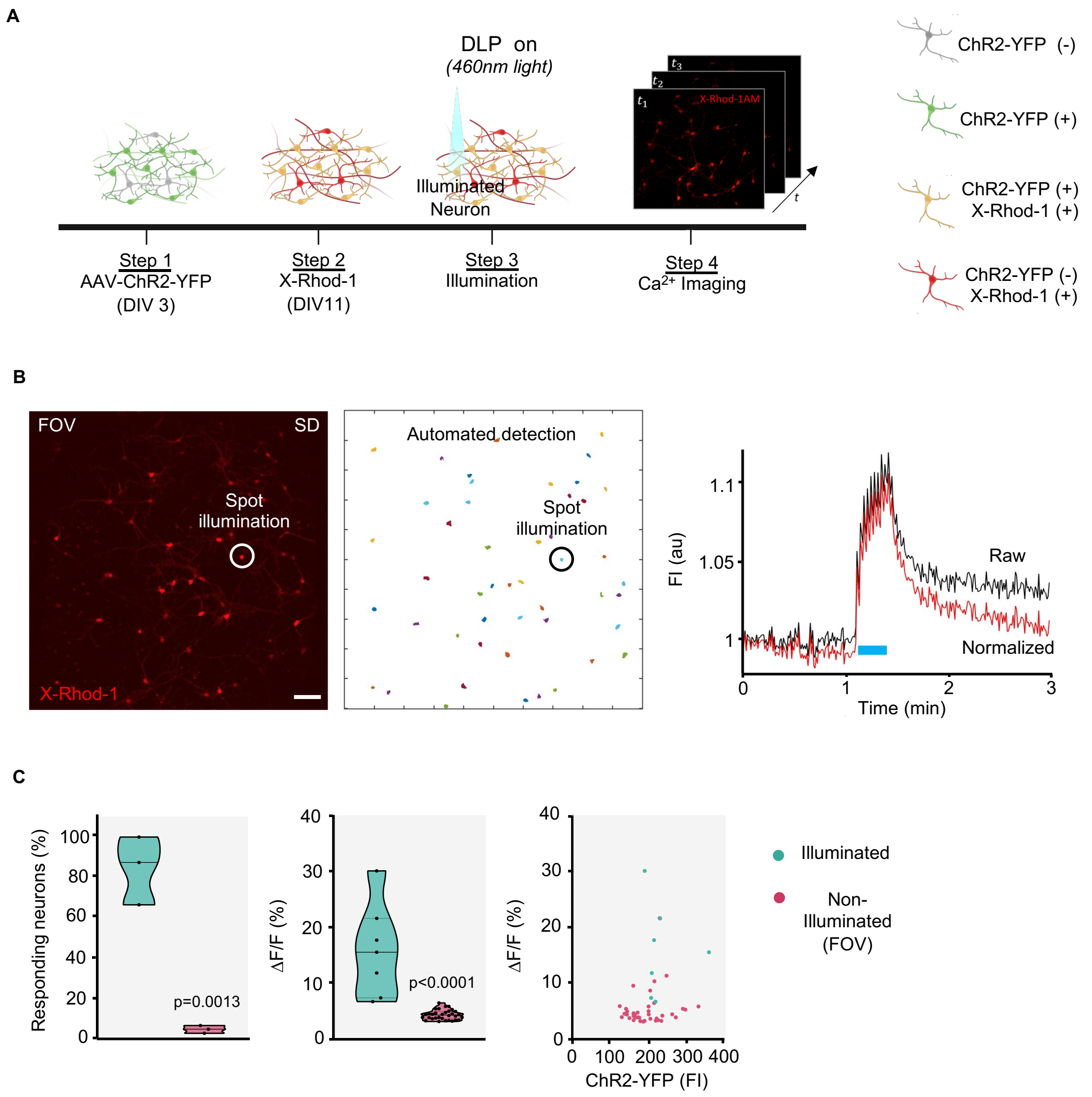
A
DLP on
(460nm light)
Illuminated
Neuron
X-Rhod-1AM
$t_1$
$t_2$
$t_3$
t
Step 1
AAV-ChR2-YFP
(DIV 3)
Step 2
X-Rhod-1
(DIV11)
Step 3
Illumination
Step 4
$Ca^{2+}$ Imaging
ChR2-YFP (-)
ChR2-YFP (+)
ChR2-YFP (+)
X-Rhod-1 (+)
ChR2-YFP (-)
X-Rhod-1 (+)
B
FOV
SD
Spot
illumination
X-Rhod-1
Automated detection
Spot
illumination
FI (au)
1.1
1.05
1
Raw
Normalized
0
1
2
3
Time (min)
C
Responding neurons (%)
100
80
60
40
20
0
p=0.0013
ΔF/F (%)
40
30
20
10
0
p<0.0001
ΔF/F (%)
40
30
20
10
0
0
100
200
300
400
ChR2-YFP (FI)
Illuminated
Non-
Illuminated
(FOV)

**Figure S3. $Ca^{2+}$ imaging in neurons individually activated by light. Related to Figure 2.**

(**A**) Schematic diagram of the experimental procedures.

(**B**) Left, SD images depicting X-Rhod-1 fluorescence signals in a representative FOV. Right, automatically detected ROIs are displayed. Spot illumination is indicated (circle). The diagrams display the X-Rhod-1 fluorescence intensity (FI) recorded from the neuronal soma of the light-stimulated (blue line) neuron as raw data (black) and following normalization (red). Scale bar: 50 μm.

(**C**) Left, quantification of illuminated and non-illuminated neurons in the FOV (n=3 experiments; Unpaired t-test). Middle, quantification of ΔF/F (%) (n=7 cells for illuminated; n=32 for non-illuminated (FOV); Unpaired t-test). Right, ΔF/F plotted against the ChR2-YFP FI (n=7 cells for illuminated; n=39 for non-illuminated (FOV)).

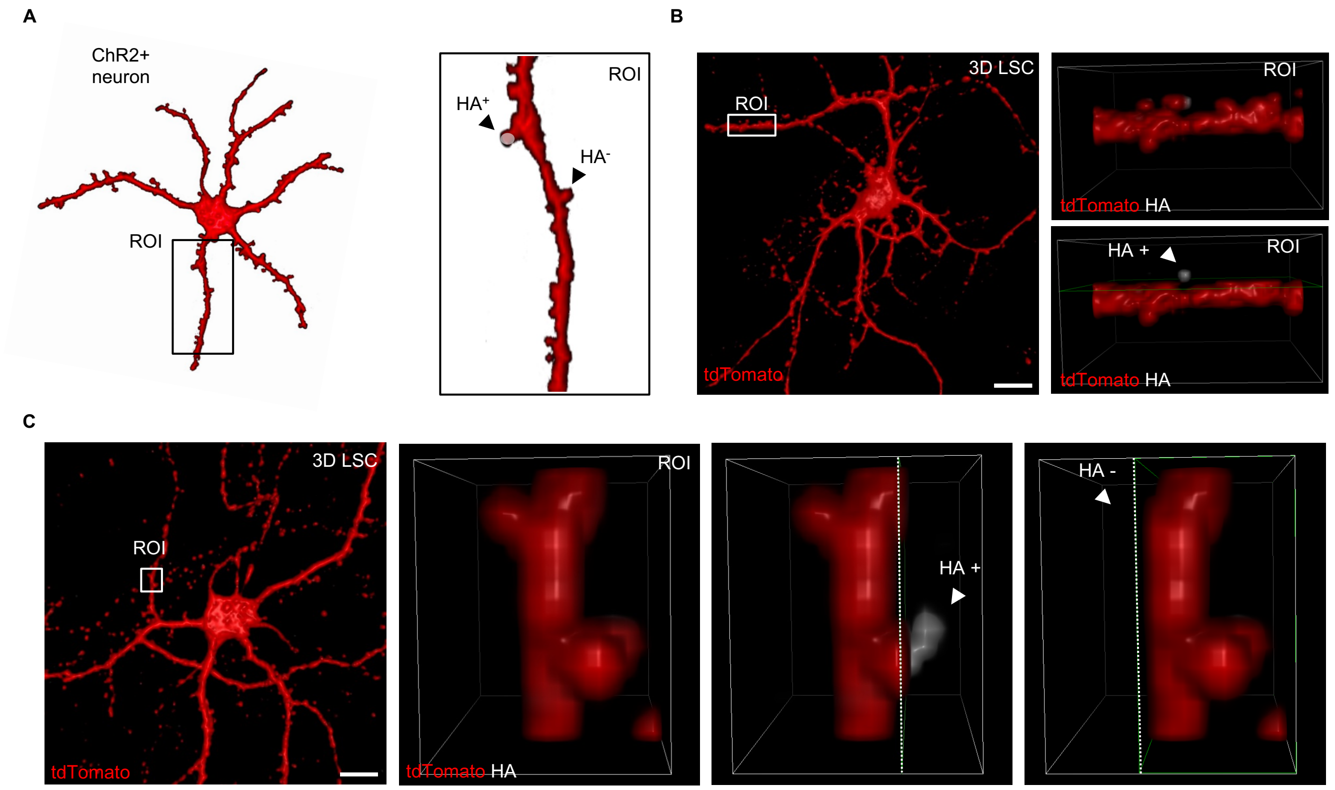

A
ChR2+
neuron
ROI
ROI
HA+
HA-
B
3D LSC
ROI
tdTomato
ROI
tdTomato HA
HA +
ROI
tdTomato HA
C
3D LSC
ROI
tdTomato
ROI
tdTomato HA
HA +
HA -

**Figure S4. HA expression in spines following light stimulation. Related to Figure 4.**

(**A**) Schematic representation of a neuron transfected with SA-tdTomato and transduced with ChR2-YFP. The ROI highlights HA+ and HA- spines.

(**B**) Left, 3D LSC image of an illuminated neuron as represented in **A**. Panels on the right depict a three-time magnification of a ROI showing a portion of a dendritic process with an HA+ spine (upper). The same ROI is presented to better visualize the HA signal (lower). Scale bar: 10 μm.

**(C)** Left, 3D LSC image of a stimulated neuron as in **A**. Panels on the right depict a ten-time magnification of a ROI (left) showing HA+ (middle) and HA- (right) spines. Scale bar: 10 μm.

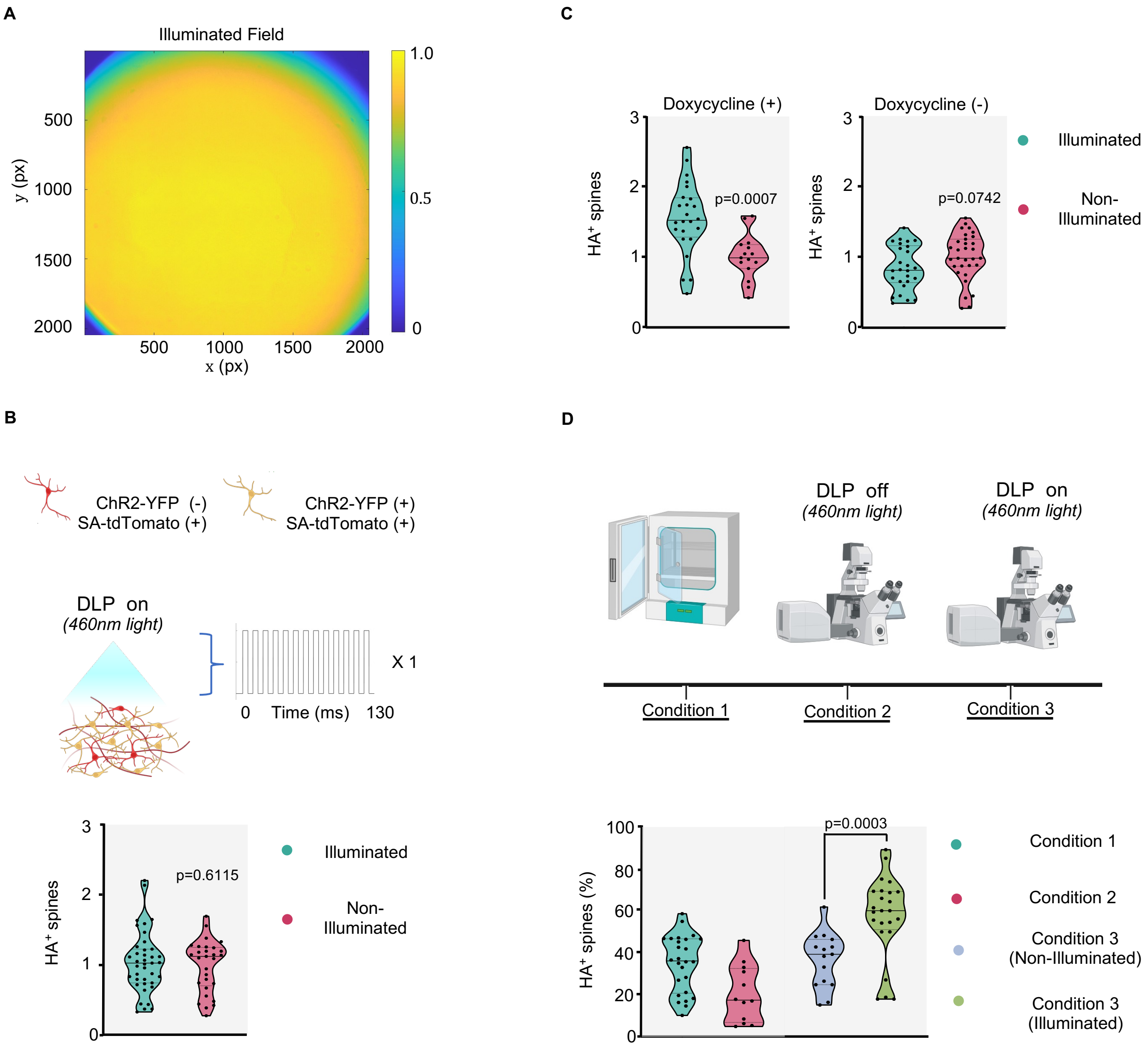

A
Illuminated Field
500
1000
1500
2000
y (px)
500
1000
1500
2000
x (px)
1.0
0.5
0
C
Doxycycline (+)
Doxycycline (-)
HA+ spines
p=0.0007
p=0.0742
Illuminated
Non-
Illuminated
B
ChR2-YFP (-)
SA-tdTomato (+)
ChR2-YFP (+)
SA-tdTomato (+)
DLP on
(460nm light)
X 1
0 Time (ms) 130
HA+ spines
p=0.6115
Illuminated
Non-
Illuminated
D
DLP off
(460nm light)
DLP on
(460nm light)
Condition 1
Condition 2
Condition 3
HA+ spines (%)
p=0.0003
Condition 1
Condition 2
Condition 3
(Non-Illuminated)
Condition 3
(Illuminated)

**Figure S5. Additional control experiments. Related to Figures 4 and Document S2.**

(**A**) Light intensity distribution on the sample plane for wide field illumination.

(**B**) Schematic diagram of the experimental procedures for single train stimulation. Violin plot illustrating the quantification of HA+ spines (%) normalized to non-illuminated neurons for 1-train illuminated and non-illuminated neurons (n=39 dendrites for illuminated; n=29 dendrites for non-illuminated; Unpaired t-test).

(**C**) Violin plots illustrating the quantification of the HA+ spines (%) for illuminated and non-illuminated neurons in the presence (left, n= 24 dendrites for illuminated; n=15 dendrites for non-illuminated; Unpaired t-test) or absence (right, n=24 dendrites for illuminated; n=15 dendrites for non-illuminated; Unpaired t-test) of doxycycline. Data are normalized to the non-illuminated neurons.

(**D**) Schematic diagram illustrating the experimental conditions (1 to 3) employed for controls. Violin plot illustrating the quantification of HA+ spines (%) in neurons kept in the incubator (condition 1), placed under the microscope without receiving DLP illumination (condition 2), and after receiving wide field DLP illumination (condition 3) (n= 26 dendrites for condition 1; n=12 dendrites for condition 2; n=15 dendrites for condition 3 non-illuminated; n=24 dendrites for condition 3 illuminated; Unpaired t-test).
